# Carbon Dots and Single-walled Carbon Nanotubes Enhances Maize Shading Stress Tolerance

**DOI:** 10.1101/2024.07.11.603111

**Authors:** Mohammad Nauman Khan, Waqar Ali, Renato Grillo, Honghong Wu, Lixiao Nie

## Abstract

Low sunlight availability/shading stress is one of the major abiotic stresses, limiting plant photosynthesis and biomass production. Maize is a C4 species and requires more sunshine for efficient photosynthesis rate. Thus, maize is a highly shade-sensitive species. We used carbon dots (CDs) and single-walled carbon nanotubes (SWCNTs) as a foliar application to enhance maize photosynthesis under no-shading and shading stress. The results revealed that under shading stress, the higher concentration of CDs and SWCNTs reduced the MDA (Malondialdehyde) content and increased the expression level of superoxide dismutase (*SOD*), peroxidase (*POD*), and catalase (*CAT*) genes. Moreover, under shading stress, CDs and SWCNTs increased the average thickness of leaf lamina, vascular bundle, mesophyll, and epidermis. CDs and SWCNTs reduced the damaging effects of shading stress on the chloroplast (Ch) formation. CDs and SWCNTs upregulated Rubisco and related genes under shading stress. The chlorophyll fluorescence parameters, including the efficiency of quantum yield of photosystem II (Fv/Fm), electron transport rate (ETR), non-photochemical quenching coefficient (NPQ), and photochemical quenching coefficient (qP) were improved with the foliar application of CDs and SWCNTs under shading stress. Higher stomatal conductance, intercellular CO_2_ concentration, transpiration, and net photosynthesis were observed in maize plants treated with CDs and SWCNTs under shading stress. The results of our study suggest that using higher concentrations of CDs and SWCNTs can enhance plant growth and photosynthesis under shading stress conditions. However, to avoid nanotoxicity, great care is recommended when selecting different concentrations of nanomaterials based on the growing conditions.

## 1. Introduction

Due to global climate change, there has been a decrease in the number of sunshine hours and solar radiation (global dimming) (Gao et al., 2020a). Urbanization and rapid economic growth have increased the emissions of air pollutants such as industrial exhausts and automobiles (Song et al., 2014). Thus, over the previous 30 years, there has been an average annual rise in haze and fog days of 16.7-33.3%, resulting in abysmal visibility (Dong et al., 2019). About 10-90 % of the incoming solar radiation has been decreased (Yang et al., 2024). The 10% decrease in incoming solar radiation might reduce crop yield by > 10% (Liu et al., 2017). Maize (*Zea mays* L.) is the third highly demanded cereal crop after wheat and rice (Erenstein et al., 2022). However, compared to wheat and rice, maize is a more versatile crop used for many purposes, including industrial products, energy, and livestock feed. Hence, maize contributes dynamically to food and nutrition security and global agri-food systems (Grote et al., 2021). Maize, a C4 species, has high photosynthetic efficiency and is highly susceptible to light limitation (Bellasio and Griffiths, 2014). Due to climatic and environmental factors (extended cloudy weather) and high plant density (self-shading with the canopy growth), maize is frequently exposed to low light stress (Gao et al., 2018). Shade stress affects maize differentially depending on the intensity and duration of shading, cultivar, and growth stage. Moreover, shade stress damages chloroplasts’ morphological and ultrastructural characteristics, inhibiting chlorophyll production and reducing the canopy’s photosynthetic capacity (Gao et al., 2020b). Thus, shading/low light limits photosynthesis, reducing biomass and yield production of maize.

Conventional farming technologies recover plant damage from abiotic and biotic stresses by one-third (Mittal et al., 2020). Therefore, traditional farming techniques fail to meet the food chain system’s increasing annual demand and supply ratio. With the advent of nanotechnology, modern agriculture is transforming into precision agriculture, maximizing output from the available resources (Mukhopadhyay, 2014). In the context of increasing plant photosynthesis, recent research has reported that applying nanomaterials (NMs) could effectively augment plant photosynthesis (Giraldo et al., 2014; Wu et al., 2017a; Zhou et al., 2021). These new-age materials might augment plant photosynthesis and feed the future world population. Different NMs have been used to tune plant photosynthesis under biotic and abiotic stresses. For example, titanium oxide nanoparticles (TiO_2_ NPs) increased photosynthesis in spinach, wheat, and tomato (Gao et al., 2008; Larue et al., 2012; Raliya et al., 2015). Cerium oxide nanoparticles (CeO_2_ NPs) were reported to increase rapeseed, *Arabidopsis*, rice, and cotton under stress conditions (Wu et al., 2017a; An et al., 2020; Liu et al., 2021; Zhou et al., 2021; Li et al., 2022a). In *Dracocephalum moldavica*, chitosan-selenium nanoparticles (Cs-Se NPs) enhanced antioxidant enzymes and photosynthetic parameters against cadmium stress (Azimi et al., 2021). Zinc oxide nanoparticles (ZnO NPs) increased wheat carbon assimilation rate and photosynthesis (Wang et al., 2020). However, the use of nanomaterials is like a double-edged sword. Nanomaterials lead to nanotoxicity and have biosafety concerns (mostly metal-based NPs). Recent studies have reported that nanotoxicity is related to the NPs characteristics (size/zeta potential/dosage/stability/coating materials), plant characteristics (plant species/growth stage), and environmental conditions (growth medium/weather conditions) (Ma et al., 2015; Yanga et al., 2017a; Khan et al., 2023). Nanotoxicity in plants has been reported to be associated with root growth inhibition, delayed plant growth and developmental stages, reduction in transpiration rate, and reduction in biomass production (Kim et al., 2012; Ma et al., 2015). Most NPs lead to toxicity at high concentrations in plants, altering their morpho-anatomical, metabolic, physiological, and genetic processes, disrupting cellular and sub-cellular organelle integrity, and reducing productivity (El-Moneim et al., 2021). NPs toxicity to plants is also due to the production of reactive oxygen species (ROS), which causes lipid peroxidation, oxidative stress, protein degradation, and DNA damage (Ma et al., 2015; Yanga et al., 2017b; Murali et al., 2022). Carbon-based nanomaterials such as carbon dots (CDs) and single-walled carbon nanotubes (SWCNTs) have been reported to have relatively less biosafety concerns and can directly interact with the plant photosynthetic system (Swift et al., 2019; Gohari et al., 2021; Li et al., 2023). The primary physiological roles of CDs in promoting crop yield and plant growth include improving nutrient uptake by the plant, stimulating root germination and seed germination, increasing biomass accumulation, improving photosynthesis, and increasing the levels of plant carbohydrates, abiotic stress, and disease resistance (Guirguis et al., 2023; Li et al., 2023). CDs were reported to enhance antioxidant enzymes, hormones, CO_2_ assimilation, movement of water and ions, chlorophyll content, electron transport rate, and Rubisco carboxylase activity, ultimately increasing the photosynthesis rate of wheat, rice, and maize (Tripathi and Sarkar, 2015; Li et al., 2021b; Wang et al., 2021a). SWCNTs are also important carbon-based NMs, playing an important role in modern agriculture (Velikova et al., 2021). SWCNTs were reported to improve antioxidants, electron transport, and photoabsorption of chloroplast (Giraldo et al., 2014). In *Chlamydomonas reinhardtii*, SWCNTs enhanced the efficiency of photosynthetic apparatus against photoinhibition (Antal et al., 2022). In rice, SWCNTs increased the expression of antioxidant enzymes, reduced oxidative damage, and enhanced photosynthesis (Zhang et al., 2017).

The present study aimed to elucidate the role of carbon-based nanomaterials such as CDs and SWCNTs in improving the photosynthesis of maize (a highly sunlight-demanding species) under no-shading shading and stress. We investigated how the foliar application of CDs and SWCNTs modulates maize biomass accumulation, oxidative damage, leaf anatomical structures, and antioxidant enzyme expression under no-shading and shading stress. Moreover, we examined the response of Rubisco (ribulose-1,5-bisphosphate carboxylase/oxygenase) and its related gene expression to the foliar application of CDs and SWCNTs under no-shading shading and stress. We also investigated the effects of CDs and SWCNTs foliar application on the efficiency of quantum yield of photosystem II (Fv/Fm), non-photochemical quenching coefficient (NPQ) and photochemical quenching coefficient (qP), electron transport rate (ETR), and net photosynthesis rate of maize crop under no-shading shading and stress. Overall, the present study was intended to enhance maize photosynthesis under low sunlight availability and also to determine the toxic and nontoxic effects of the foliar application of CDs and SWCTNs under no-shading shading and stress.

## 2. Materials and Methods

### 2.1 Nanoparticles description

Single-walled carbon nanotubes (Cat #XFS19) and carbon dots (Cat #XF253) were obtained from Jiangsu XFNANO Materials Tech Co., Ltd. in PR, China. Using an FEI Talos microscope running at 300 kV, transmission electron microscopy (TEM) pictures were captured. The dispersion of CDs and SWCNTs in DI water was assessed for zeta potential using a dynamic light scattering apparatus (Malvern Zetasizer, Nano). The CDs and SWCNTs were sonicated for 30 minutes at 37 kHz using an ultrasonic cleaner to encourage uniform suspension and reduce aggregation.

### 2.2 Experimental details and nanoparticle treatment

A pot experiment was conducted at the glasshouse of Hainan University, Haikou, PR, China, to investigate the effects of CDs and SWCNTs foliar application on the maize growth performance under the no-shading and shading stress. According to our former experimental procedures, we used shading nets to impose 60% shading stress (Song et al., 2022). This study used the maize cultivar “Zhengdan958 (obtained from Shandong Shouhe Seed Industry Co., Ltd)” as an experimental material. Pots with a height of 21.3 cm, a lower diameter of 18.3 cm, and an upper diameter of 23.3 cm were filled with 4 kg soil (originated from Lingao, Danzhou, Hainan Province, China, sun-dried and sieved 2 mm). Compound fertilizer ∼6.67 g (N: P: K, 15: 15: 15) was uniformly mixed with each pot’s soil. Before sowing, the maize seeds were evenly spread out on wet germination papers in a growth chamber after being soaked in DI water for 16 hours. The germinated seedlings were planted in the pots immediately after their radicle length measured 2 mm. Shading nets were positioned 60 centimeters above the potted plants following a week of the seedlings’ growth in the pots. The maize seedlings (25 and 32 DAS) were treated with CDs and SWCNTs via foliar spray (∼5 mL/plant) at 0.1, 0.2, and 0.3 mg/L. These concentration ranges were determined by reviewing earlier research [39, 42–44] that documented the beneficial effects of CDs and SWCNTs. DI water was sprayed for non-nanoparticle (NNP) as a control treatment. Using an electronic balance, the fresh weight of the maize seedlings was noted at 60 DAS. When necessary, the irrigation, pesticides, and insecticides were applied in the same way to every treatment.

### 2.3 Maize leaf anatomy and cellular ultrastructure analysis

Leaf samples of maize plants treated with CDs, SWCNTs, and NNP were collected to study the differences in the anatomy of the various tissues. Utilizing techniques outlined in the earlier study [45], 3 mm^2^ leaf samples were sliced with a sharp knife. After 2h in a safranin O staining solution, the samples were exposed to 50%, 70%, and 80% ethanol grades. The specimens were then dehydrated with anhydrous ethanol and fixed in a solid green plant solution for 20 min. The samples spent 5 min in xylene. Using an upright optical microscope, the images of the leaf tissue were captured (Nikon Eclipse E100).

Fresh leaf samples from the plants treated with CDs, SWCNTs, and NNP control were collected following the previously described protocols [42]. The leaf samples were cut into 1 mm^3^ blocks with a sharp blade. The 1 mm^3^ leaf blocks were put immediately into 2 mL tubes with a fixative solution (Servicebio, G1102). The samples were preserved by first being fixed at ambient temperature until they sank to the bottom, and then they were kept at 4 °C. Subsequently, the samples underwent three 15-minute rinses with 0.1 M PB (pH 7.4). The specimen samples were polymerized for 48 hours at 65 °C and implanted (Acetone: EMBed 812) at 37 °C overnight. The tissues were adhered to on 150 mesh cuprum grids with formvar film after thinning the resin blocks to 60–80 nm using the ultra-microtome. The ultrathin sections were cleaned three times with ultra-pure water, three times with 70% ethanol, and eight minutes of staining with 2% uranium acetate. After inserting the cuprum grids into the grid board, they were left to cure overnight at room temperature. The TEM images were captured using a Hitachi (HT7800) transmission electron microscope.

### 2.4 Measurement of leaf photosynthesis-related traits

On a bright day, from 10:00 AM to 2:00 PM, the portable photosynthetic equipment (Li-6400, Li-COR Inc.) was utilized to monitor the intercellular CO_2_ concentration, transpiration rate, stomatal conductance, and photosynthesis rate. The characteristics associated with photosynthesis were recorded using the fully expanded third maize flag leaf. The following settings were made for the portable photosynthetic system: air humidity at 70%, light intensity at 1000 µmol m^-2^ s^-1^, leaf temperature at 23 °C, and CO_2_ at 400 µmol mol^-1^. A PAM-2500 chlorophyll fluorometer (Heinz Walz GmbH, Germany) was used to estimate the chlorophyll fluorescence parameters. To determine chlorophyll fluorescence parameters, the third fully developed maize flag leaf was chosen at 8:00 PM in a dark environment. 200 µmol m^-2^ s^-1^ was the actinic light intensity, while 1200 µmol m^-2^ s^-1^ was the saturation light intensity. The study assessed the following parameters: electron transport rate (ETR), non-photochemical quenching coefficient (NPQ), photochemical quenching coefficient (qP), and maximum quantum yield of PSII (Fv/Fm) (Wang et al., 2015).

### 2.5 Determination of membrane damage and quantification of antioxidants

Malondialdehyde (MDA) content was measured to estimate oxidative damage [49]. 1 mL of 5% (w/v) trichloroacetic acid was used to homogenize 0.1 g of fresh leaf material before centrifuging at 10,000 g for 10 min. 0.4 mL of the supernatant was added with 0.4 mL of 5% TCA that contained 0.67% (w/v) thiobarbituric acid. For about 15 min, the samples were submerged in a 95°C hot water bath. The samples were centrifuged at 10,000g for 5 min after cooling on ice following the hot water bath. On the UV 1800PC, AOE, Shanghai, China spectrophotometer, readings were taken at 450, 532, and 600 nm for the final calculation of MDA content (Khan et al., 2019). By homogenizing a 0.1 g fresh leaf sample with 1 mL 50 mM PBS (pH 7.8), the antioxidant enzyme activities, including CAT (EC 1.11.1.6), SOD (EC 1.15.1.1), and POD (EC 1.1.1.1.7), were determined. After obtaining the supernatant, it was centrifuged for 20 minutes at 4°C at 12,000 × g. To determine CAT [50], SOD [51], and POD [52], respectively, ODs were measured at 240 nm, 560 nm, and 470 nm using a spectrophotometer (UV 1800PC, AOE, Shanghai, China).

### 2.6 Quantification of Rubisco enzyme activity and content

To estimate the amount of ribulose-1,5-bisphosphate carboxylase/oxygenase (Rubisco) and determine its enzymatic activity, Jiangsu Meibiao Biological Technology Co., Ltd. ELISA kits (MB-10606B) were used. To ascertain the concentration and activity of Rubisco, 0.1 g fresh leaf samples were used. The maize leaf samples’ Rubisco content and activity were assessed using the Rubisco ELISA Kit and calibration standards. Samples and calibration standards were run alongside to create a standard curve. Then, the Rubisco enzyme activity and content in the samples were determined by comparing the sample ODs with the standard curve.

### 2.7 RNA extraction, cDNA preparation, and qPCR experiment

Trizol reagent (Invitrogen, Santa Clara, CA, USA) was used to extract RNA from fresh leaf samples. SuperScript III reverse transcriptase (Invitrogen, Grand Island, NY, USA) performed reverse transcription operations following the manufacturer’s instructions. To synthesize cDNA, oligo(dT)20 primers were used. Using a Bio-rad Mx3000P qPCR system and SYBR Premix Ex Taq II (Tli RNaseH Plus) (TaKaRa Biotech. Co.), qRT-PCR analyses were carried out. The primer design was done using Primer3 software. In this work, the reference gene was 18s (XR_004853753.1). Amplification technique [53] was used, which included 30 seconds of denaturation at 95°C, 40 cycles of 95°C for 5 seconds, 60°C for 30 seconds, and 72°C for 30 seconds, and a final extension step at the end. The melting curve was obtained by heating the amplicon from 65°C to 95°C at a rate of 0.5°C per second. The qRT-PCR detection was carried out in biological duplicates three times. The 2^−ΔΔCt^ technique was utilized to evaluate the relative expression levels.

### 2.8 Statistical analysis

Using Statistix 9.0, all recorded indices were subjected to a one-way ANOVA analysis. The data shown is the average of four individual replications with standard error (± SE) values. At the 0.05 probability levels, the significant difference between the treatment means was differentiated using the least significance difference (LSD) test. The figures were constructed with Origin 2021 (Origin Lab. Crop.).

## 3. Results

### 3.1 Characterization of CDs and SWCNTs NPs

The size of the CDs was 2–5 nm, according to the TEM (Fig. 1A). The TEM image of SWCNTs reveals that the length was 1-3 µm, the outer diameter (OD) was 1-2 nm, and the inner diameter (ID) was 0.8-1.6 nm (Fig. 1B). The zeta potential of CDs and SWCNTs, according to analysis from the Malvern, Zetasizer, Nano, was ∼-5mV and ∼-30 mV, respectively (Fig. 1C).

### 3.2 Foliar application of CDs and SWCNTs has ameliorative effects on maize plant height and biomass production under shading stress

The results from the screening experiment showed that the foliar application of CDs and SWCNTs at the concentrations of 0.1, 0.2, and 0.3 mg/L has different effects on the plant height and biomass production of maize (Figure 1D-E and Additional File Supplementary Figure S1). Briefly, under no-shading stress, the foliar application of CDs and SWCNTs at higher concentrations (0.3 mg/L) reduced the plant height and biomass production of maize compared to non-nanoparticle treatment (NNP). However, under shading stress, the higher concentration of CDs and SWCNTs (0.3 mg/L) increased the plant height and biomass production of maize compared to the NNP treatment (Figure 1E). Following the objective of the present research of tuning maize photosynthesis and biomass production of maize under shading stress, we selected the higher concentration of CDs and SWCNTs to explore the mechanism behind shade tolerance or toxicity. We observed 18.78% and 22.22% decrease in maize plant height with the foliar application of CDs and SWCNTs (0.3 mg/L), respectively, compared to NNP under no-shading growing conditions. While under shading stress, the foliar application of CDs and SWCNTs increased maize plant height to 21.83% and 17.48, respectively, compared to NNP (Figure 1D). Similar trend was recorded for the maize biomass production with the foliar application of CDs and SWCNTs under no-shading and shading stress (Figure 1E). The results of the present reveal that under no-shading, the foliar of CDs and SWCNTs at higher concentrations (0.3 mg/L) harmed maize growth and biomass production. However, the higher concentration of CDs and SWCNTs improved the maize growth and biomass production under shading stress.

**Figure 1.**
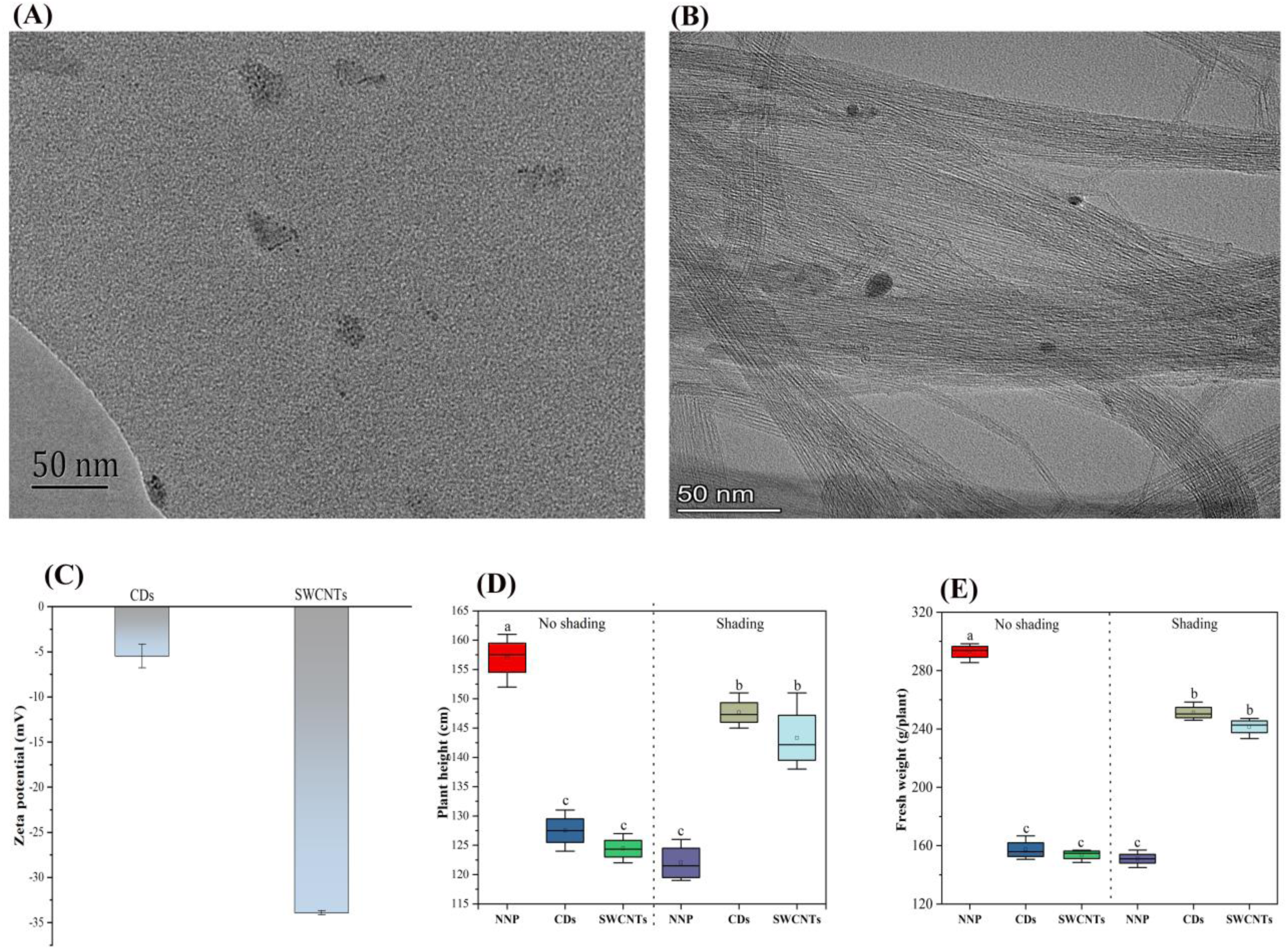
The characterization and effects of CDs and SWCNTs on maize growth under no-shading and shading stress. **(A)** TEM image of CDs (Scale bar: 50 nm), **(B)** TEM image of SWCNTs (Scale bar: 50 nm), **(C)** zeta potential of CDs and SWCNTs, **(D)** effect of CDs and SWCNTs foliar application on maize plant height under no-shading and shading stress, and **(E)** effect of CDs and SWCNTs foliar application on maize biomass production under no-shading and shading stress. NNP; non-nanoparticle treatment, CDs; carbon dots (0.3 mg/L), SWCNTs; single-walled carbon nanotubes (0.3 mg/L). The means were separated using the LSD test at the 0.05 probability levels, and the significance among the treatment means is represented with different lowercase alphabets. Error bars on boxes represent the standard error of four individual replications.

### 3.3 CDs and SWCNTs decreased the oxidative damage under shading stress

The foliar application of CDs and SWCNTs increased MDA content by 237.07% and 226.79%, respectively, compared to NNP under no shading stress (Figure 2A). Whereas, under shading stress, CDs and SWCNTs reduced the MDA content by 26.48% and 28.76, respectively, compared to NNP treatment. NNP, CDs, and SWCNTs did not significantly affect the POD activity under no-shading stress (Figure 2B). However, a 276.40% and 263.61% increase in POD activity was recorded for the maize plants treated with CDs and SWCNTs, respectively, compared to NNP under shading stress. The highest SOD activity was recorded for CDs (281.04%) and SWCNTs (278.26%) under the shading stress (Figure 2C). Similarly, the highest CAT activity was observed for the maize plants treated with CDs (189.51%) and SWCNTs (175.02%) under the shading stress (Figure 3D). Moreover, the foliar application of CDs and SWCNTs downregulated *POD*, *SOD*, and *CAT* genes expression under no-shading stress, compared to NNP (Figure 2E-G). However, the *POD*, *SOD*, and *CAT* genes were upregulated by the foliar application of CDs and SWCNTs under the shading stress, compared to NNP (Figure 2E-G).

**Figure 2.**
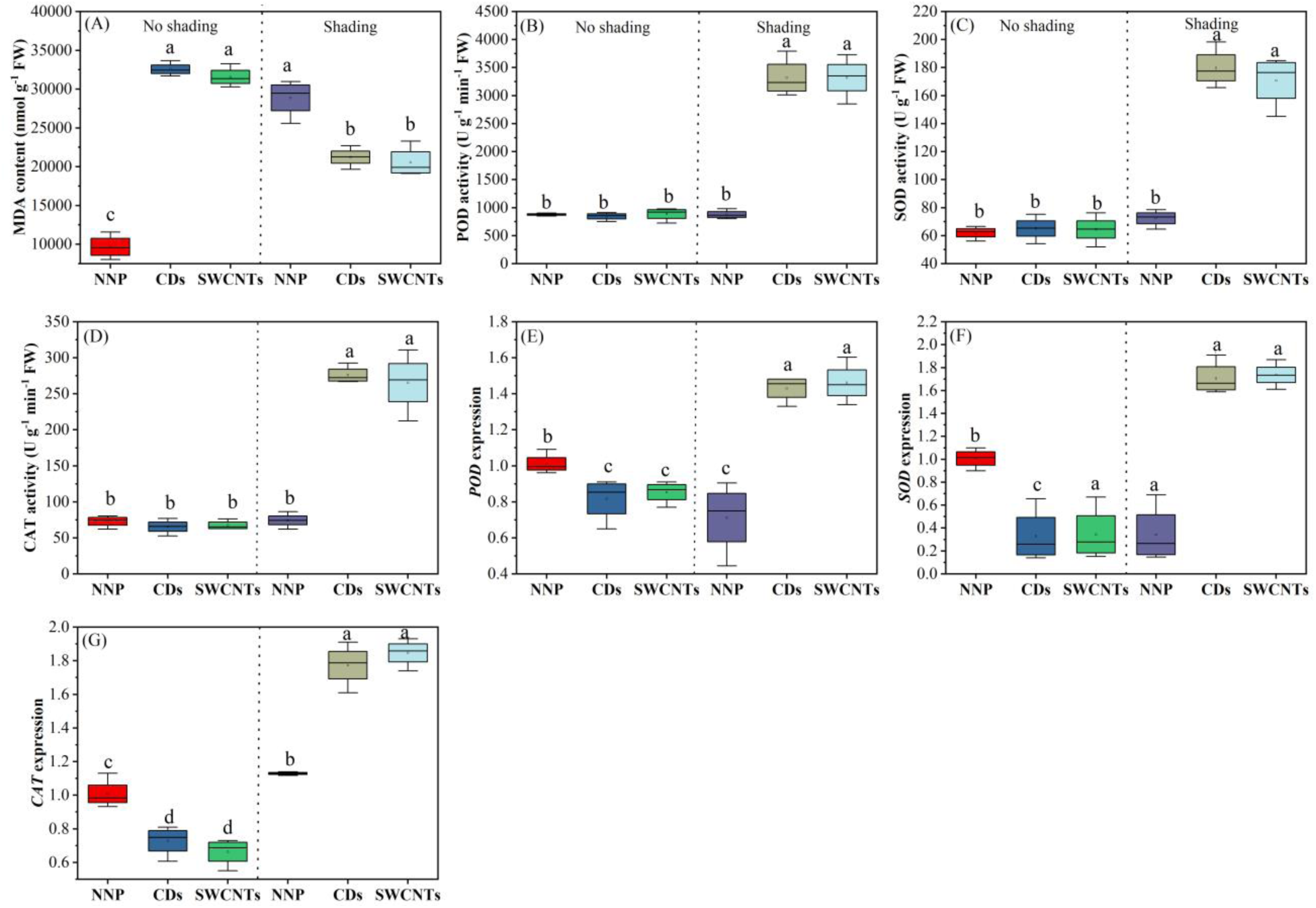
The regulation of MDA content and antioxidant enzymes by the foliar application of CDs and SWCNTs under no-shading and shading stress. **(A)** response of MDA content to foliar application of CDs and SWCNTs under no-shading and shading stress **(B, C, and D)**, response of POD, SOD, and CAT activities to foliar application of CDs and SWCNTs under no-shading and shading stress, and **(E, F, G)** regulation of response of *POD*, *SOD*, and *CAT* genes expression by foliar application of CDs and SWCNTs under no-shading and shading stress. NNP; non-nanoparticle treatment, CDs; carbon dots (0.3 mg/L), SWCNTs; single-walled carbon nanotubes (0.3 mg/L). The means are separated using the LSD test at the 0.05 probability levels, and the significance among the treatment means is represented with different lowercase alphabets. Error bars on boxes represent the standard error of four individual replications.

**Figure 3.**
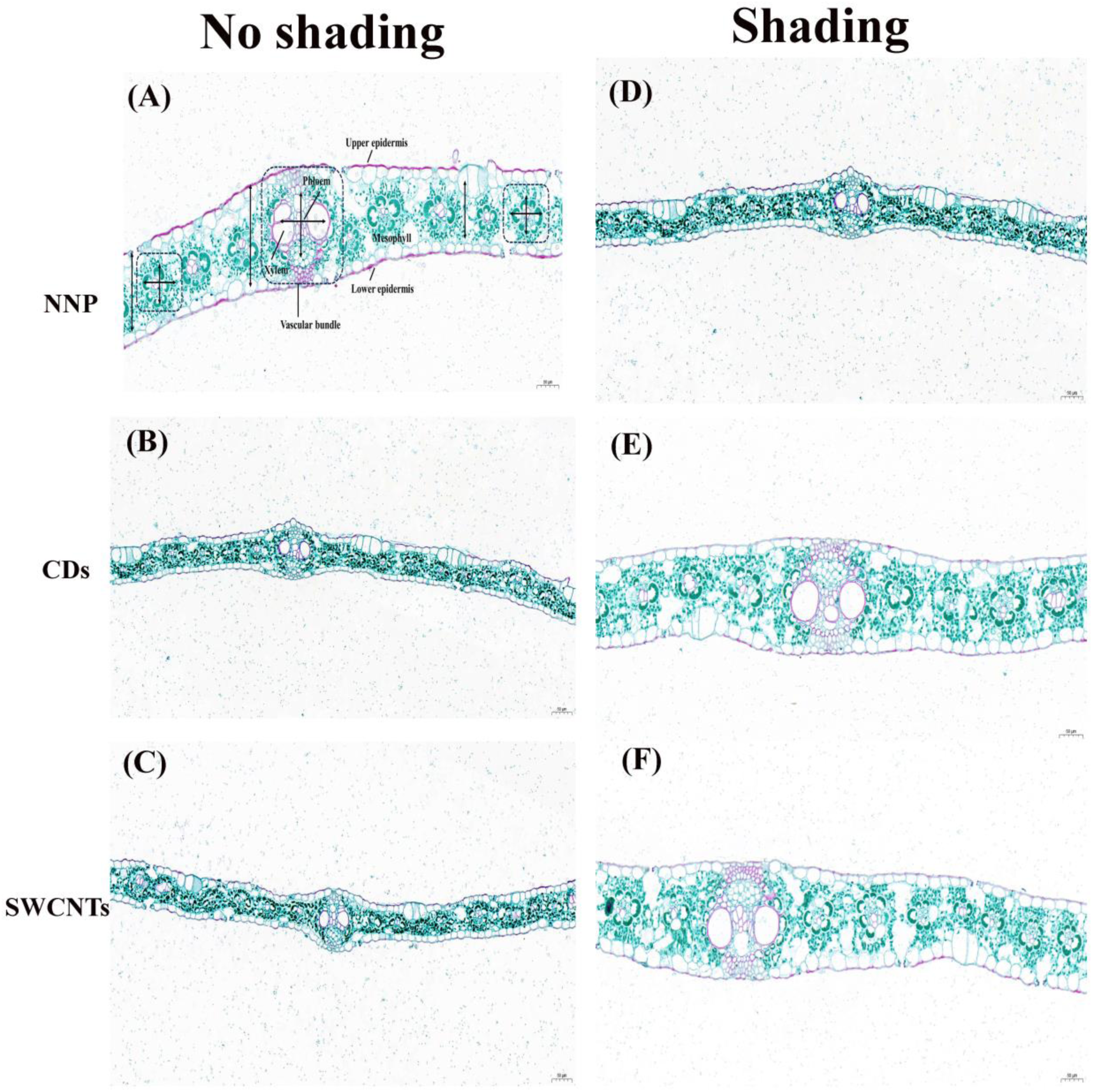
The effects of foliar application of CDs and SWCNTs on maize leaf anatomical structures under no-shading and shading stress. **(A, B, and C)** effect of foliar application of CDs and SWCNTs on the average thickness of leaf lamina, vascular bundle, mesophyll, and epidermis under no-shading, and **(D, E, and F)**, the effect of foliar application of CDs and SWCNTs on the average thickness of leaf lamina, vascular bundle, mesophyll, and epidermis under shading stress. NNP; non-nanoparticle treatment, CDs; carbon dots (0.3 mg/L), SWCNTs; single-walled carbon nanotubes (0.3 mg/L). The images were taken through an upright optical microscope (Nikon Eclipse E100) for at least four individual samples—scale bar=50 µm.

### 3.4 Maize leaf anatomy and cellular ultrastructure were ameliorated under shading stress with foliar application of CDs and SWCNTs

Under no-shading, compared to NNP, CDs and SWCNTs reduced the average leaf lamina thickness by 54.30% and 55.28%, respectively (Table 1). Under shading stress, CDs and SWCNTs increased the average leaf lamina thickness by 71.99% and 75.29%, respectively (Table 1). Compared to NPP, a reduction in average vascular bundle thickness was recorded for CDs (67.94%) and SWCNTs (84.06%) under no-shading stress (Table 1 and Figure 3). However, a 51.90% and 60.80% increase in the average vascular bundle thickness was recorded for CDs and SWCNTs under shading stress, compared to NNP under no-shading stress (Table 1 and Figure 3). CDs and SWCNTs reduced the average mesophyll thickness by 82.15% and 81.97%, respectively, compared to NNP under no-shading (Table 1 and Figure 3). While CDs and SWCNTs increased the average mesophyll thickness by 233.35% and 244.79%, respectively, compared to NNP under shading stress (Table 1 and Figure 3). Similarly, under no-shading, compared to NNP, CDs and SWCNTs reduced the average epidermis thickness by 59.58% and 60.46%, respectively (Table 1 and Figure 3). However, CDs and SWCNTs increased the average epidermis thickness by 72.10% and 65.02%, respectively, compared to NNP under shading stress (Table 1 and Figure 3).

**Table 1.**
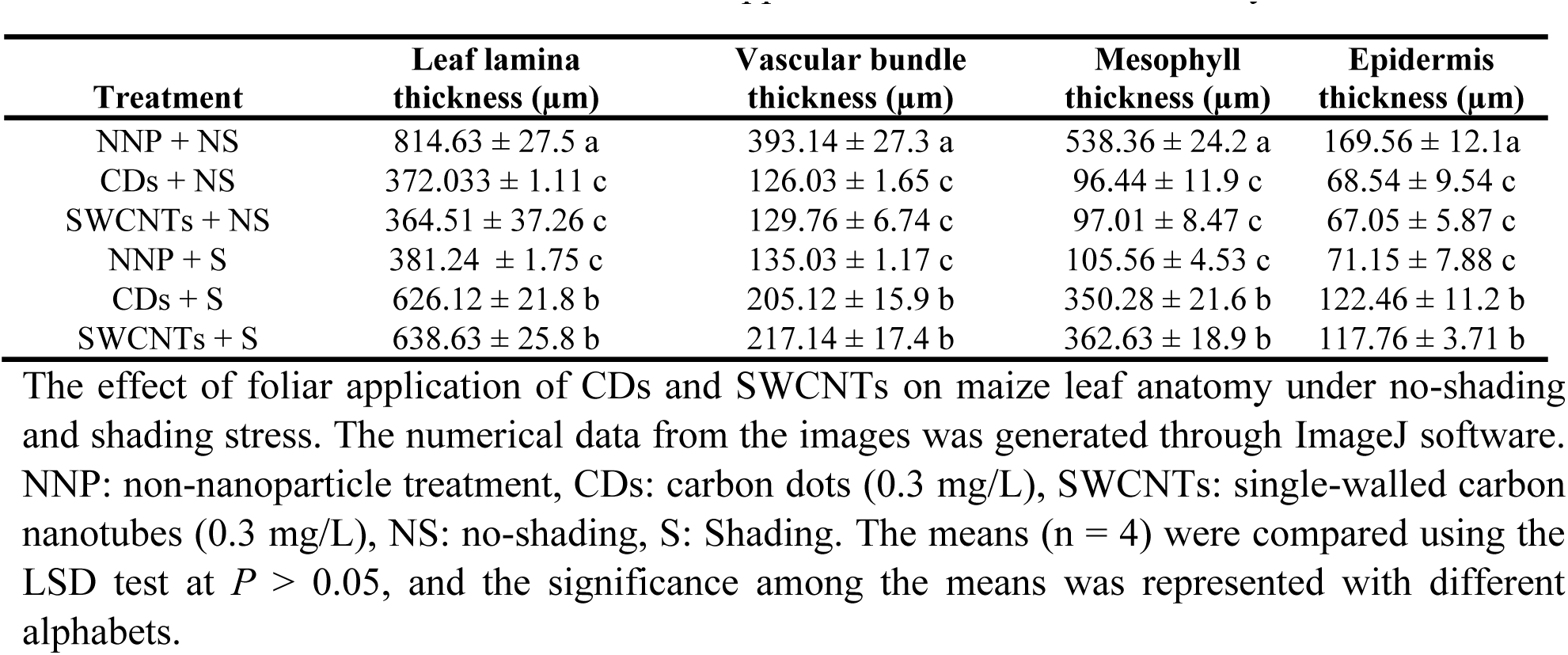
Effect of CDs and SWCNTs foliar application on maize leaf anatomy.

| Treatment | Leaf lamina thickness ($\mu\text{m}$) | Vascular bundle thickness ($\mu\text{m}$) | Mesophyll thickness ($\mu\text{m}$) | Epidermis thickness ($\mu\text{m}$) |
| --- | --- | --- | --- | --- |
| NNP + NS | 814.63 ± 27.5 a | 393.14 ± 27.3 a | 538.36 ± 24.2 a | 169.56 ± 12.1a |
| CDs + NS | 372.033 ± 1.11 c | 126.03 ± 1.65 c | 96.44 ± 11.9 c | 68.54 ± 9.54 c |
| SWCNTs + NS | 364.51 ± 37.26 c | 129.76 ± 6.74 c | 97.01 ± 8.47 c | 67.05 ± 5.87 c |
| NNP + S | 381.24 ± 1.75 c | 135.03 ± 1.17 c | 105.56 ± 4.53 c | 71.15 ± 7.88 c |
| CDs + S | 626.12 ± 21.8 b | 205.12 ± 15.9 b | 350.28 ± 21.6 b | 122.46 ± 11.2 b |
| SWCNTs + S | 638.63 ± 25.8 b | 217.14 ± 17.4 b | 362.63 ± 18.9 b | 117.76 ± 3.71 b |

The effect of foliar application of CDs and SWCNTs on maize leaf anatomy under no-shading and shading stress. The numerical data from the images was generated through ImageJ software. NNP: non-nanoparticle treatment, CDs: carbon dots (0.3 mg/L), SWCNTs: single-walled carbon nanotubes (0.3 mg/L), NS: no-shading, S: Shading. The means (n = 4) were compared using the LSD test at $P > 0.05$ , and the significance among the means was represented with different alphabets.

The cellular ultrastructure analysis showed that compared to NNP, CDs and SWCNTs reduced the cell size of maize leaf under no-shading (Figure 4A-C). Similarly, smaller chloroplasts with autopepsia were observed under no-shading in the maize leaf treated with CDs and SWCNTs. Moreover, loose cell walls with no clear boundaries were observed in CDs and SWCNTs treated maize leaf cells under no-shading, compared to NNP. In the NPP treatment under shading stress, smaller cells, smaller chloroplasts with autopepsia, and loose cell walls with no precise edges were observed (Figure 4D). However, under shading stress, CDs and SWCNTs increased the cell size, chloroplast size with no autopepsia, and well-organized cell walls with precise edges around the cells were overserved (Figure 4E-F).

**Figure 4.**
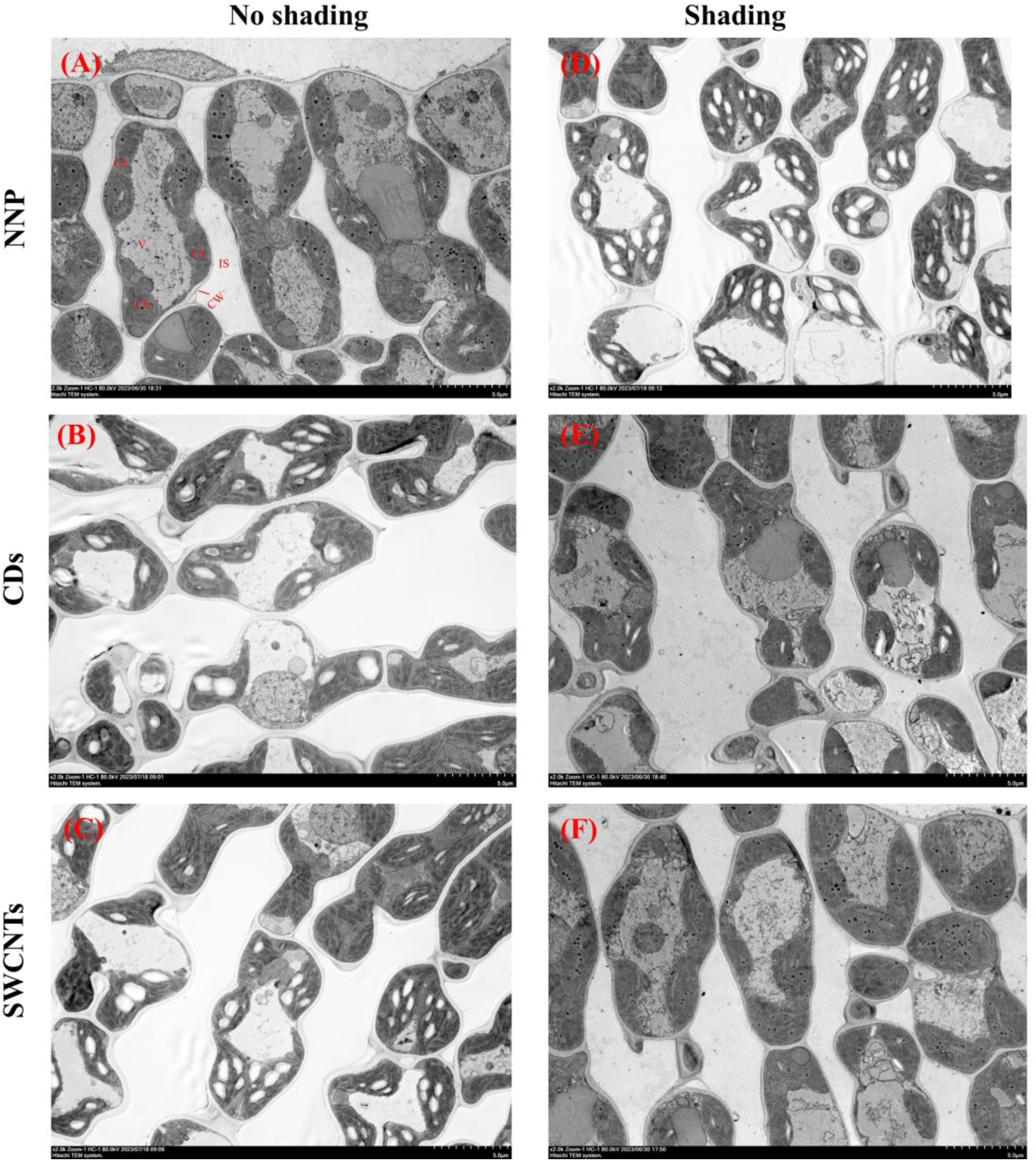
The effects of foliar application of CDs and SWCNTs on maize leaf cells ultrastructure under no-shading and shading stress. **(A, B, and C)** effect of foliar application of CDs and SWCNTs on cell size, cell wall formation, and chloroplast formation under no-shading, and **(D, E, and F)**, the effect of foliar application of CDs and SWCNTs on cell size, cell wall formation, and chloroplast formation under shading stress. NNP; non-nanoparticle treatment, CDs; carbon dots (0.3 mg/L), SWCNTs; single-walled carbon nanotubes (0.3 mg/L). The TEM images were taken through a transmission electron microscope (Hitachi, HT7800) for at least four individual samples. Scale bar =5 µm.

### 3.5 CDs and SWCNTs increased Rubisco content and activity under shading stress

Compared to NNP under no-shading, the foliar application of CDs and SWCNTs decreased the Rubisco activity and content by 58.86%, 50.27%, and 62.53%, 50.80%, respectively (Figure 5A-B). However, compared to NNP under shading, the foliar application of CDs and SWCNTs increased the Rubisco activity and content by 51.74%, 50.16%, and 51.27.53, 53.65%, respectively (Figure 5A-B). Under no-shading stress, CDs and SWCNTs downregulated the expression of *rbcL*, *rbcS*, *RAF1*, and *Rca* genes, compared to NNP (Figure 5C-F). Nonetheless, CDs and SWCNTs upregulated the *rbcL*, *rbcS*, and *Rca* expression under shading stress, compared to NNP (Figure 5C, D, and F). The *RAF1* gene was downregulated under shading and no-shading stress for all the treatments (Figure 5E).

**Figure 5.**
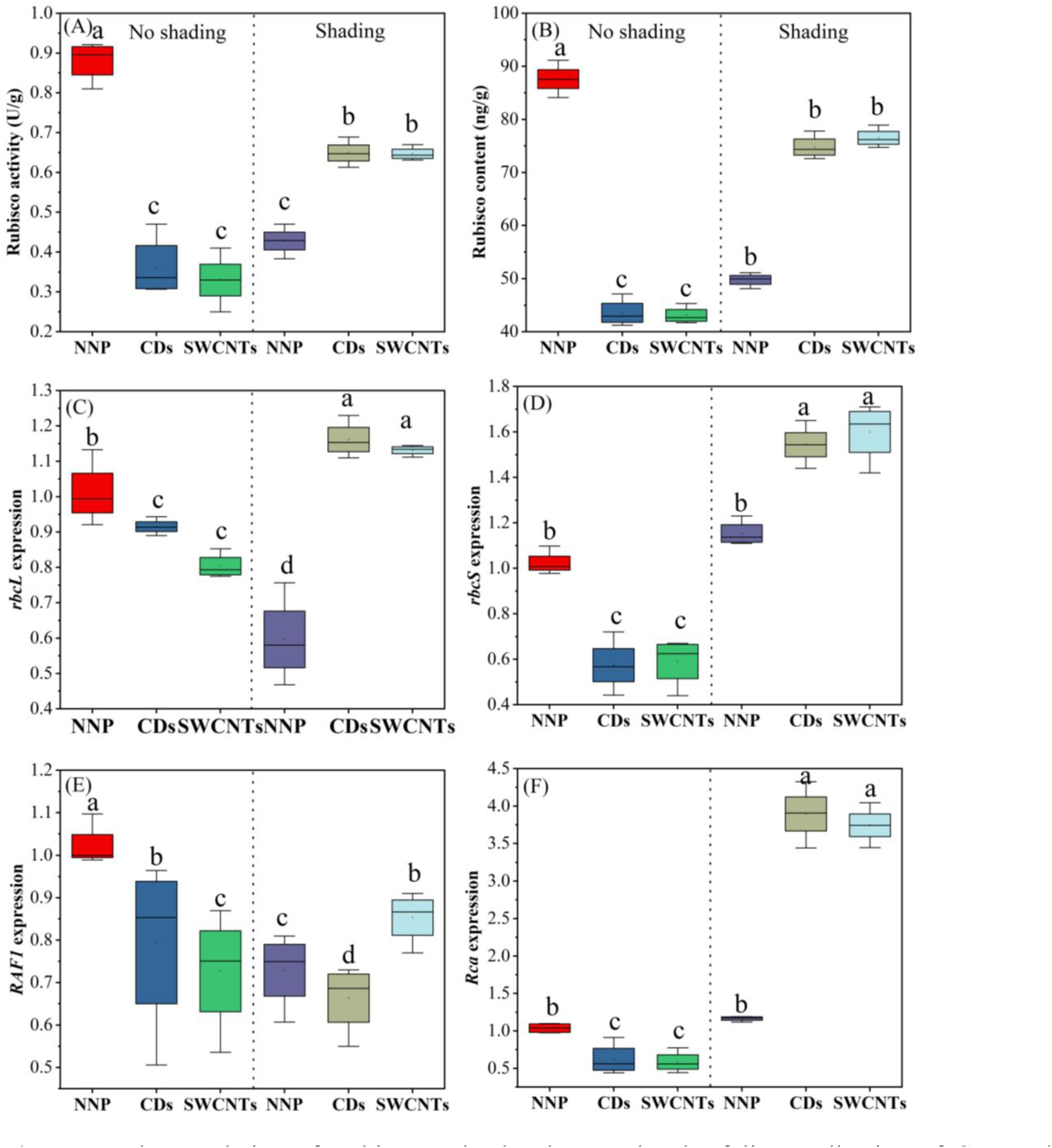
The regulation of Rubisco and related genes by the foliar application of CDs and SWCNTs under no-shading and shading stress. **(A, B)** response of Rubisco enzyme activity and content to foliar application of CDs and SWCNTs under no-shading and shading stress, and **(C, D, E, and F)**, regulation of *rbcL*, *rbcS*, *RAF1*, and *Rca* genes expression by foliar application of CDs and SWCNTs under no-shading and shading stress. NNP; non-nanoparticle treatment, CDs; carbon dots (0.3 mg/L), SWCNTs; single-walled carbon nanotubes (0.3 mg/L). The means are separated using the LSD test at the 0.05 probability levels, and the significance among the treatment means is represented with different lowercase alphabets. Error bars on boxes represent the standard error of four individual replications.

### 3.6 Photosynthesis and its related components enhanced under shading stress by CDs and SWCNTs

The maximum quantum yield of PSII (Fv/Fm) was significantly reduced by 68.58% and 69.71% by the foliar application of CDs and SWCNTs, respectively, under no-shading stress, compared NNP (Figure 6A). Under shading stress, a 198.46% and 129.16% increase in maximum quantum yield of PSII (Fv/Fm) was observed for CDs and SWCNTs treated maize plants, respectively, compared to NNP. CDs (61.80%) and SWCNTs (58.38%) significantly reduced electron transport rate (ETR) compared to NPP under no-shading stress (Figure 6B). There was a significant increase up to 52.05% and 46.48% in ETR under shading stress for CDs and SWCNTs, respectively, compared to NNP. Under no-shading, CDs and SWCNTs reduced the non-photochemical quenching coefficient (NPQ) by 68.69% and 63.35%, respectively, compared to NNP (Figure 6C). While under shading stress compared with NNP, CDs and SWCNTs increased NPQ by 143.92% and 131.07%. Under no-shading stress compared with the NNP, the photochemical quenching coefficient (qP) was decreased by 43.43% and 34.33% by the foliar application of CDs and SWCNTs, respectively (Figure 6D). Meanwhile, qP was increased by 334.75% and 424.81% by the foliar application of CDs and SWCNTs, respectively, compared with NNP (Figure 6D).

**Figure 6.**
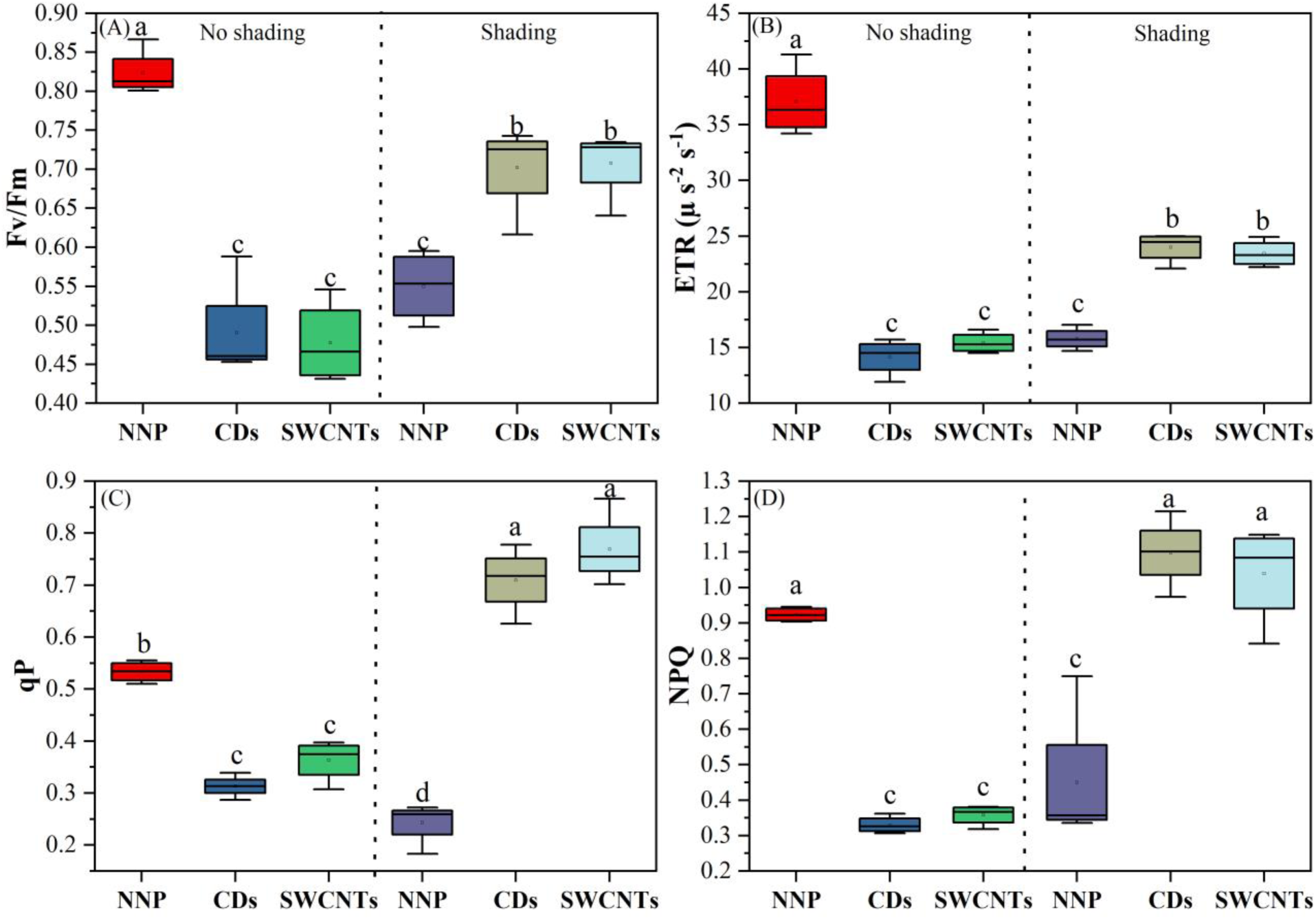
The effect of foliar application of CDs and SWCNTs on the maize chlorophyll fluorescence parameters under no-shading and shading stress. **(A)** response of Fv/Fm to foliar application of CDs and SWCNTs under no-shading and shading stress, **(B)** response of ETR to foliar application of CDs and SWCNTs under no-shading and shading stress, **(C)** response of qP to foliar application of CDs and SWCNTs under no-shading and shading stress, and **(D)** response of NPQ to foliar application of CDs and SWCNTs under no-shading and shading stress. NNP; non-nanoparticle treatment, CDs; carbon dots (0.3 mg/L), SWCNTs; single-walled carbon nanotubes (0.3 mg/L). The means are separated using the LSD test at the 0.05 probability levels, and the significance among the treatment means is represented with different lowercase alphabets. Error bars on boxes represent the standard error of four individual replications.

**Figure 6.**
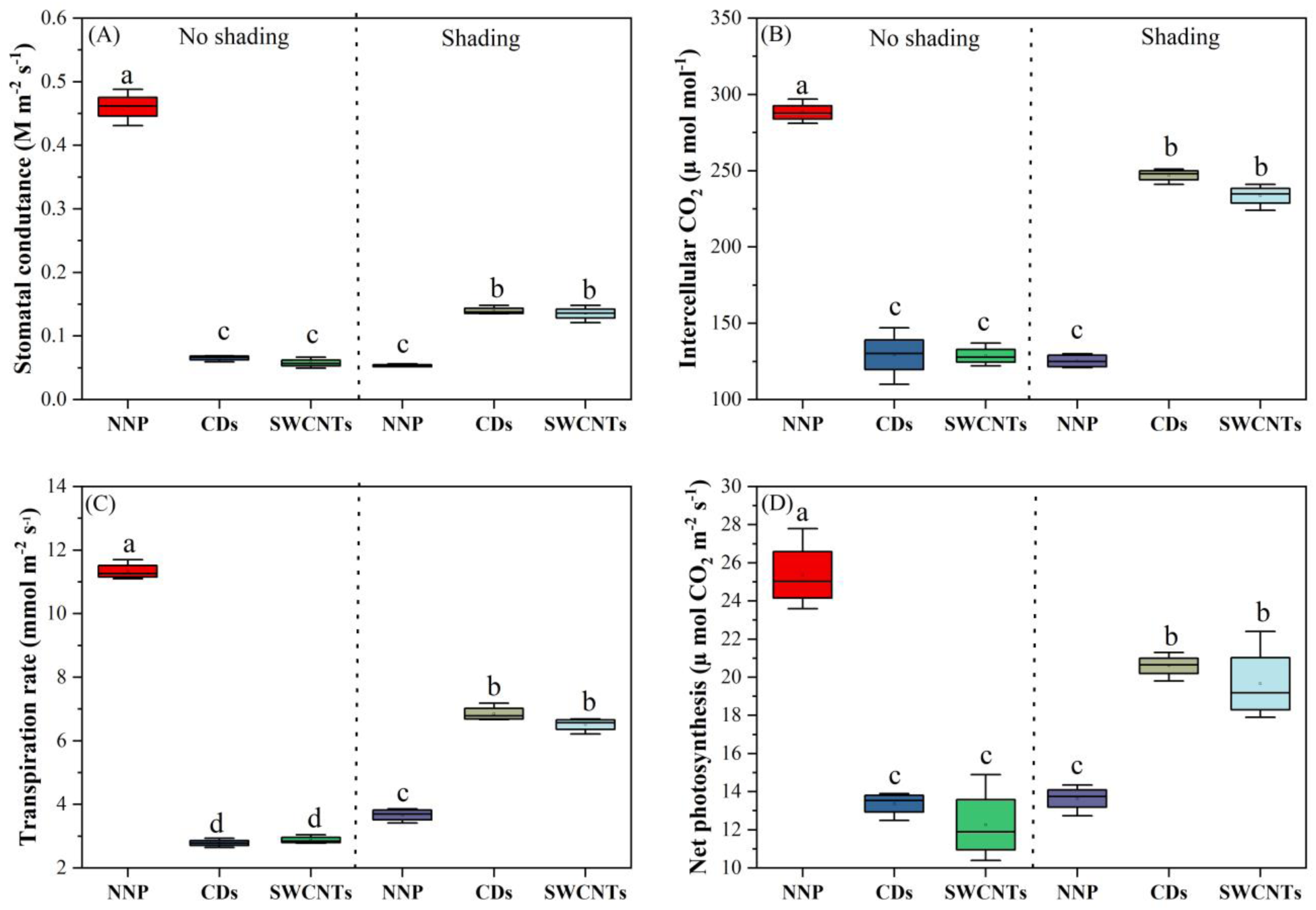
The effect of foliar application of CDs and SWCNTs on the maize chlorophyll fluorescence parameters under no-shading and shading stress. **(A)** response of Fv/Fm to foliar application of CDs and SWCNTs under no-shading and shading stress, **(B)** response of ETR to foliar application of CDs and SWCNTs under no-shading and shading stress, **(C)** response of qP to foliar application of CDs and SWCNTs under no-shading and shading stress, and **(D)** response of NPQ to foliar application of CDs and SWCNTs under no-shading and shading stress. NNP; non-nanoparticle treatment, CDs; carbon dots (0.3 mg/L), SWCNTs; single-walled carbon nanotubes (0.3 mg/L). The means are separated using the LSD test at the 0.05 probability levels, and the significance among the treatment means is represented with different lowercase alphabets. Error bars on boxes represent the standard error of four individual replications.

The stomatal conductance (gs) was significantly reduced up to 85.50% and 87.53% by the foliar application of CDs and SWCNTs, respectively, compared to NNP under no-shading stress (Figure 7A). Under shading stress compared to NNP, CDs and SWCNTs increased gs by 162.86% and 154.70%, respectively. The intercellular CO_2_ concentration (ci) was reduced by the foliar application of CDs (55.14%) and SWCNTs (55.37%), compared to NPP under no-shading stress (Figure 7B). However, compared to NNP under shading stress, ci was increased by CDs (97/20%) and SWCNTs (86.56%). The transpiration rate (E) was reduced by the foliar application of CDs (75.44%) and SWCNTs (75.61%), compared to NNP under no-shading stress (Figure 7C). CDs (86.90%) and SWCNTs (77.54%) increased the transpiration rate, compared to NNP under shading stress. The maize photosynthesis rate (Pn) was significantly reduced by the foliar application of CDs (47.30%) and SWCNTs (51.65%), compared to NNP under no-shading stress (Figure 7D). However, the Pn rate was significantly enhanced by the foliar application of CDs (51.02%) and SWCNTs (44.18%), compared to NNP under shading stress.

## 4. Discussion

Maize is a C4 species that demands high sunlight for robust growth and development (Wang et al., 2014). Besides the reduced sunshine hours and solar radiation due to climate change, cloudy weather, and self-shading during the later growth stage, maize is exposed to low light/shading stress (Wang et al., 2014; Gao et al., 2018). Shading stress inhibits maize photosynthesis, decreasing biomass and yield production (Gao et al., 2020a). Similarly, the results of our study revealed that shading stress reduced the maize plant height (22.29%, Figure 1D) and biomass accumulation (48.42%, Figure 1E). We used CDs and SWCNT NPs as a foliar application to enhance maize shade tolerance. In the present study, we studied the effects of CDs and SWCNTs on maize physiological responses under shading stress. We observed that under shading stress, plant height and biomass production of maize were increased due to the foliar application of CDs and SWCNTs (Figure 1D-E and Additional File Figure S1). Thus, the findings of our study suggest that the effects of CDs and SWCNTs are not only related to the concentrations of the NPs but also the growing conditions. To elucidate further the mechanism behind the dual effects of NPs on maize growth under no-shading and shading stress, we selected the higher concentration (0.3 mg/L) of CDs and SWCNTs.

The balance between oxidative damage and the regulation of antioxidant enzymes controls plant growth and development (Considine et al., 2015; Li et al., 2022b; Kumar et al., 2024). Compared to NNP, we found a significant increase in MDA content in CDs and SWCNTs treated maize plants, indicating the toxicity of the NPs under no-shading stress (Figure 2A). However, CDs and SWCNTs reduced the MDA content under shading stress, reflecting the ameliorative effects of NPs under shading stress (Figure 2A). It is well documented that NPs may generate reactive oxygen species (ROS) depending on concentration and growing conditions, causing nanotoxicity (Ray et al., 2009; Considine et al., 2015; Khan et al., 2023). In the present study the possible reason behind the oxidative damage under no-shading stress for the higher concentrations of CDs and SWCNTs could be the lower activities of antioxidant enzymes and the downregulation of related genes expression, including *POD* (Figures 2B and E), *SOD* (Figures 2C and F), and *CAT* (Figures 2D and G). The nanotoxicity of carbon-based NPs due to oxidative stress generation in various plant species has been reported in many recent reports (Verma et al., 2019; Jiang et al., 2020; Li et al., 2020). Moreover, under shading stress, the higher CDs and SWCNTs increased the enzymatic activities and upregulated the gene expression of *POD* (Figures 2B and E), *SOD* (Figures 2C and F), and *CAT* (Figures 2D and G). It shows that triggering antioxidant enzymes may be one of the mechanisms of improving maize growth and biomass production under shading stress. Up to date, the primary and commonly shared mechanism behind NPs-induced stress tolerance in plant species is associated with the scavenging of ROS through upregulating antioxidant enzyme system (Wu et al., 2017a; Newkirk et al., 2018; Wu et al., 2018; Swift et al., 2019).

Generally, shade stress results in thinner and smaller leaves (Wu et al., 2017c). Due to the lower dry mass per unit area, the thinner leaves have a higher chance of intercepting light. However, the thinner leaves have less capacity for CO_2_ dissolution and transportation (Gong et al., 2015; Wu et al., 2017c). Similarly, in the present study, while comparing the NNP treatments under no-shading and shading stress, we observed that shading stress reduced the average thickness of the maize leaf lamina (53.18%), vascular bundle thickness (83.41%), mesophyll thickness (80.48%), and epidermis thickness (58.05%) (Table 1 and Figure 3). Moreover, under no-shading, CDs and SWCNTs significantly reduced the average leaf lamina, vascular bundle, mesophyll, and epidermis thickness compared to NNP (Table 1 and Figure 3). This could possibly be explained as the higher concentrations of CDs and SWCNTs increased the oxidative damage under no-shading stress and thus reduced the maize leaf growth. However, under shading stress, CDs and SWCNTs at higher concertation upregulated the antioxidant enzyme system, increasing the average leaf lamina, vascular bundle, mesophyll, and epidermis thickness compared to NNP (Table 1 and Figure 3). Furthermore, the cellular ultrastructure analysis showed that shading stress reduced the cell and chloroplast size and damaged cell wall formation of maize leaf (Figure 5A and D). These results are consistent with the findings of previous studies (Gonzalez et al., 2012; Gong et al., 2014). The effect of CDs and SWCNTs under no-shading and shading stress, less cell wall damage and less reduction in cell and chloroplast size were observed under shading stress. This could be ascribed to the higher oxidative damage under no-shading stress due to higher concentration of CDs and SWCNTs while lower oxidative damage under shading stress (Figure 2). Under oxidative stress, higher antioxidant enzyme actives have been reported to protect cellular ultrastructure in different plant species (Du, 2015; Hazeem et al., 2019; Nauman et al., 2019; Velikova et al., 2021).

Rubisco activity is a key metabolic mechanism that modulates photosynthetic capability to adapt to incoming light under shade (Taylor et al., 2022). Low light/ shading stress downregulates the gene expression of Rubisco (Sun et al., 2014). The results of our study showed that compared to no-shading, shading stress downregulated the Rubisco activity (Figure 4A), content (Figure 4B), and gene expression (*rbcL*, *rbcS*, *RAF1*, and *Rca* genes, Figure 4C-F). The lower Rubisco activity and content under no-shading stress can be ascribed to the damaging effects of the foliar application of CDs and SWCNTs, while the maize plants showed downregulation of Rubisco-related genes, including *rbcL*, *rbcS*, *RAF1*, and *Rca* (Figure 4C-F). However, CDs and SWCNTs enhanced Rubisco activity, content, and related gene expression under shading stress (Figure 4A-F). The upregulation of the Rubisco enzyme due to NPs application under different stresses has been reported in various plant species (Gao et al., 2008; Giraldo et al., 2014; Wu et al., 2017a; Li et al., 2018a; Tighe-Neira et al., 2018; Li et al., 2021b). Under shading stress, CDs and SWCNTs enhanced the efficiency of PSII (Figure 5A), ETR (Figure 5B), qP (Figure 5C), and NPQ (Figure 5D). It is evident from the ultrastructure analysis that CDs and SWCNTs ameliorated the damaging effects of shading on chloroplast formation (Figure 4A-F). Thus, under shading stress with less oxidative damage (Figure 2A) and chloroplast preservation, CDs and SWCNTs sustained higher Fv/Fm, ETR, qP, and NPQ. Previously, it was reported that CDs worked as a light converter from UV radiation to PAR and accelerated ETR in PSII (Li et al., 2021a; Li et al., 2021b; Wang et al., 2021b). Similarly, SWCNTs were also reported to enhance PSII, ETR, qP, and NPQ in plant species under different stress conditions (Zhang et al., 2017; Petrova et al., 2021; Wu et al., 2023). Shading stress decreased stomatal conductance, intercellular CO_2_ concertation, transpiration rate, and photosynthesis rate (Figure 7A-D). It is well known that under limited sunlight, the reduction in plant photosynthesis is related to lower stomatal conductance, transpiration rate, and lower uptake of CO_2_ from the atmosphere (Wu et al., 2017b; Dong et al., 2019; WANG et al., 2021). Depending on the growing conditions (no-shading vs shading), SWCNTs significantly reduced the leaf anatomical structures under no-shading while enhancing the leaf anatomical structures under shading stress (Figure 3A-F). Under shading stress, CDs and SWCNTs increased the average thickness of the vascular bundle, mesophyll cells, and epidermis layers, sustaining higher leaf lamina thickness. Thicker leaves have more efficiency of light absorption, CO_2_ uptake and contain more chloroplasts for the synthesis of chlorophyll content (Jiang et al., 2011; Gonzalez et al., 2012; Gong et al., 2014; Gong et al., 2022). Therefore, compared to NNP treatment under shading stress, CDs and SWCNTs had higher stomatal conductance, intercellular CO_2_ concentration, transpiration, and photosynthesis rates (Figure 7A-D). Not surprisingly, NPs have been reported to have ameliorative effects on plant photosynthesis under hostile environmental conditions (Perreault et al., 2012; Wang et al., 2016; Zhang et al., 2017; Zhang et al., 2018; Velikova et al., 2021; Guirguis et al., 2023).

## 5. Conclusion

The use of nanomaterials could be an effective way to augment maize photosynthesis under shading stress. Under shading stress, the higher concentrations of CDs and SWCNTs increased the maize growth and photosynthesis. The mechanism behind CDs and SWCNTs-improved maize biomass and photosynthesis included the upregulation of antioxidant enzymes genes, improving leaf anatomical structures, persevering chloroplast, upregulation of Rubisco-related genes, increasing chlorophyll fluorescence, and photosynthesis-related traits. The findings of the present study suggest that depending on the growing environment, the optimum concentration of CDs and SWCNTs can be an effective way to increase plant photosynthesis under shading stress. However, field experiments are recommended to explore further the increase in crop yields with the use of nanomaterials such as CDs and SWCNTs under the shading.

## Authors’ contributions

MNK, LN, HW: Conceptualization, MNK, WA, Conducting experiment and providing support, MNK, NL, HW: Writing-review, MNK, WA, RG: Figures and tables construction, MNK, LN, HW, RG: Visualization, supervision, funding acquisition.

## Acknowledgements

Authors are grateful to all laboratory members of the Sanya Nanfan Research Institute of Hainan University for their support.

